# Fungi exposed to chronic nitrogen enrichment are less able to decay leaf litter

**DOI:** 10.1101/067306

**Authors:** Linda T.A. van Diepen, Serita D. Frey, Elizabeth A. Landis, Eric W. Morrison, Anne Pringle

## Abstract

Saprotrophic fungi are the primary decomposers of plant litter in temperate forests, and their activity is critical for carbon (C) and nitrogen (N) cycling. Simulated atmospheric N deposition is associated with reduced fungal biomass, shifts in fungal community structure, slowed litter decay, and soil C accumulation. Although rarely studied, N deposition may also result in novel selective pressures on fungi, affecting evolutionary trajectories. To directly test if long-term N enrichment reshapes fungal behaviors, we isolated decomposer fungi from a longterm (28 year) N addition experiment and used a common garden approach to compare growth rates and decay abilities of isolates from control and N amended plots. Both growth and decay were significantly altered by long-term exposure to N enrichment. Changes in growth rates were idiosyncratic, but litter decay by N isolates was generally lower compared to control isolates of the same species, a response not readily reversed when N isolates were grown in control (low N) environments. Changes in fungal behaviors accompany and perhaps drive previously observed N-induced shifts in fungal diversity, community composition, and litter decay dynamics.

## INTRODUCTION

The fungi are a megadiverse group of microbial species that play critical roles in biogeochemical cycles. Anthropogenically-mediated environmental changes are a serious threat to biological diversity and may result in novel selective pressures (Sala et al. 2000), but whether or how fungi are adapting to global change is largely unknown. Vulnerable fungi may move or go extinct in response to environmental stresses. Shifts in fungal phenology in response to climate change are documented (Boddy et al. 2014), as are fungal range expansions (Wolfe et al. 2012). However, whether fungi are prone to extinction remains an open question, largely because conservation efforts rarely target fungi (Heilmann-Clausen et al. 2015). Species may also be tolerant of changes, either because the stresses are irrelevant to the species in question, or because the species can alter aspects of behavior or life history in response to changing environments (West-Eberhard 1989).

Fungi are not static entities, and fungal lineages can evolve on very short time scales. For example, individuals exposed to novel environments in the laboratory diverge and evolve over weeks or months (Schoustra et al. 2005; Leu and Murray 2006; Dettman et al. 2007, 2008). However, there is little research on the evolution of fungi in global change contexts, even though evolutionary dynamics may accompany or drive shifts in fungal diversity that occur in response to environmental change (Jaenike 1991; Avis et al. 2008; Bärlocher et al. 2008). As fungal lineages evolve in response to global change, the physiology and function of these species may change, and the evolutionary trajectories taken by individual lineages may aggregate to influence ecosystem scale processes.

Fungi are the primary drivers of decomposition in temperate forest systems (Gadd 2006), and altered rates of decomposition will directly influence future trajectories of climate change (Kirschbaum 1995; Ise and Moorcroft 2006). Since saprotrophic fungi are the primary producers of the extracellular enzymes involved in cellulose and lignin (lignocellulose) decay, it may be especially critical to understand the evolution of saprotrophic fungi in global change contexts. Global change experiments, in which the abiotic environment is manipulated to simulate future scenarios of a particular (or multiple) global change driver(s) in the field, offer a unique opportunity to explore changes in species’ behaviors relevant to ecosystem-scale processes, although these kinds of experiments are rarely used for this purpose (Bataillon et al. 2016). In this study, we use a long-term (28 year) simulated nitrogen (N) deposition experiment—the Chronic Nitrogen Amendment Study (CNAS) located at the Harvard Forest Long-Term Ecological Research (LTER) site in Petersham, MA, USA—to explore fungal behaviors in contexts of altered soil N availability. Although not originally designed as an artificial selection experiment, a continuous and consistent exposure to elevated N within treatment plots has likely acted as a novel selective pressure on fungi. Previous research from the CNAS shows that longterm N enrichment reduces fungal biomass, causes changes in fungal diversity and community composition (Frey et al. 2014; Morrison et al. 2016), slows plant litter decay (Magill and Aber 1998), and results in a significant accumulation of soil organic matter (Frey et al. 2014). However, whether changes in fungal behaviors accompany or drive shifts in decomposition dynamics remains untested.

Here we investigated whether the saprotrophic fungi that grow on plant litter of N-enriched plots behave differently from the same species found in control plots. Have these isolates evolved, compared to isolates of the same species from control plots, and does that evolution impact decomposition? We first isolated fungi from decomposing plant litter in control and N-enriched plots and next used a common garden design to study the behaviors of these individuals in their respective home and away environments. To measure potential changes in fungal behaviors, we first measured mycelial growth rate, often assumed to be an aspect of fitness (Pringle and Taylor 2002) and next measured the ability of isolates to decay plant litter. We compared isolates from the N treatments with isolates of the same species from control plots, growing all isolates in identical laboratory environments (the same “gardens”) where N availability was varied to reflect field conditions in the different N treatments at CNAS. Our working assumption was that if fungal isolates from N-enriched plots grew similarly to those from control plots when grown in a common environment, then isolates from treatment plots have not evolved. By contrast, if we found differences—if isolates from the N treatment plots showed consistent differences in growth and litter decay compared to control isolates—then data would suggest that the lineages isolated from N treatment plots are distinct and that treatment isolates have evolved to grow in the novel environments of the experimental plots.

## METHODS

### Fungal cultures

Fungi were isolated from decomposing leaf litter collected from each of three N addition treatments within the CNAS which was established in 1988 to examine the effects of simulated N deposition on ecosystem dynamics (Aber et al. 1993). The site is located in a mixed hardwood forest dominated by black and red oak *(Quercus velutina* and *Q. rubra)*. The experiment consists of three 30 × 30 m plots which receive either ambient N deposition (control, N0) or an additional 50 (N50) or 150 (N150) kg N ha^−1^ yr^−1^. The CNAS has resulted in a wealth of data (Aber et al. 1993; Magill et al. 2004; Frey et al. 2004; Frey et al. 2014; Morrison et al. 2016), and while it has been criticized for pseudo-replication, it remains one of the longest running N addition experiments in existence. For this reason, it is a particularly appropriate choice for experiments exploring fungal evolution where the continuous and consistent exposure to elevated N has likely acted as a novel selective pressure on the fungi growing within the plots. While soil fungi occasionally grow very large (Smith et al. 1992) and it is possible that a single individual could grow between plots, the plots are large enough to encompass the entire canopies and root structures of multiple individual trees (>100 in each plot; Fig. S1) and would, by extension, house the fungal communities associated with these trees, especially saprotrophic taxa housed within individual decomposing leaves and litter patches.

Litter samples for culturing fungi were obtained from a litterbag experiment described in van Diepen et al. (2015). The culture media used to isolate fungi and the protocol used to identify them are detailed in the Supplementary Information. Briefly, we targeted those fungal species that were isolated from both the control and N treatment plots. To minimize the probability of using the same individual twice, we never used more than one isolate of a species from the same litterbag and made sure that isolates of the same species came from different litterbags collected at least 5 m apart. Approximately 1500 isolates were cultured, and in this isolate collection, more than 100 different morphologies were distinguishable. A subset of 60 morphologies was found in both the control and at least one N treatment, and following identification using the internal transcribed spacer (ITS) barcode, the different morphologies grouped into a collection of 41 different species. The majority of species were from the phylum Ascomycota or Zygomycota, and 21 species (51%) matched an operational taxonomic unit (OTU) found in a high-throughput sequencing dataset (Morrison et al. in prep) obtained from the same litterbags. Nineteen of these 21 species had either representative isolates from all three field treatments or representatives from the control and one of the N treatments. Subsequent experiments targeted ten ascomycetes (Supplementary Table S1). All were 1) identified and grouped using the ITS barcode, 2) found in the high-throughput sequencing dataset (providing a context for future work) and 3) found in the control and at least one N treatment plot. In the sequencing dataset, the relative abundances of the ten taxa ranged from 0.01 to 1.2% (Table S1), typical for saprotrophic taxa at our site (Morrison et al. 2016). Collectively, the relative abundance of these taxa was 1.4% in the control treatment, increasing to 1.9 and 3.9% in the N50 and N150 treatments, respectively. We also included one basidiomycete in this study, although basidiomycetes were harder to culture as they were mostly outgrown by ascomycetes on all culture media. To facilitate communication, we assigned a provisional name to each group of isolates based on the best match to a voucher species within NCBI, with a minimum requirement of 97% sequence similarity and a bit score of 800 (Table S1). From this point forward, we refer to each of our species using its provisional generic epithet. Seven of the 11 selected taxa were available as multiple biological replicates (more than one isolate was found in each plot). Eight of the species were cultured from all three N treatments, but two *(Discosia* and *Phacidium)* were only isolated from the control and N50 treatments and one *(Irpex)* was only isolated from the control and N150 treatments.

### Fungal growth and litter decay

To measure potential changes in the physiology of isolates, we compared the behaviors of isolates from the two N treatments (N isolates) with isolates of the same species collected from the control treatment (control isolates). We use the term “isolate origin” to indicate the field treatment (N0, N50, or N150) from which each culture was isolated and “laboratory environment” to indicate the N treatment simulated by our common garden experiment (described below). Isolates from the control treatment were grown in their home environment, as well as N50 and N150 (away) environments, and isolates from the N treatments were grown in the control (away) environment and their home (N50 or N150) environment.

Mycelial growth rates are often used as a measure of fungal fitness (Pringle and Taylor 2002). Here we compared mycelial growth rates among isolates of the same species by growing them at 25°C in darkness in ‘race tubes’ (sensu White and Woodward 1995) created by filling serological pipettes with a horizontal strip of culture medium: 1.5% potato dextrose agar dissolved in half-strength Hoagland nutrient solution supplemented with inorganic N at levels representative of the available N (NH_4_ + NO_3_) in the organic soil horizon of each respective field treatment (Supplementary Fig. S2). Growth was measured daily until each isolate reached the end of its race tube when growth rate was calculated in mm day^−1^. We ran three laboratory replicates per biological isolate in each environment, and whenever possible, growth rates were calculated for the multiple biological replicates per species isolated from each field treatment.

To test the ability of the fungal isolates to decompose plant litter, we used a common garden approach where fungi isolated from both control and N enriched field plots were grown in the laboratory in their respective home and away environments. Petri dishes were filled with 30 ml of culture medium (1.5% agar dissolved in half-strength Hoagland nutrient solution) which was then covered with ~0.75 g of sterile oak litter (Fig. S3a). The underlying medium mimicked the organic (O_e_/O_a_) horizon in the field plots, with N levels representative of available N (NH_4_ + NO_3_) within each of the three field treatments (N0, N50, or N150), while the oak litter mimicked the O_i_ (litter) layer. Fresh oak litter was collected from each treatment plot using litter traps. Collected oak litter was air dried at room temperature and the leaves cut into ~2 x 2 cm pieces after stem removal. Before use, litter was oven-dried (60°C for 48 hours) and subsequently sterilized (three rounds of autoclaving at 121°C). Because these experiments were extraordinarily time intensive, we made the deliberate choice to include more species with a single biological replicate each, rather than fewer species with more biological replicates. We manipulated one biological replicate per species per treatment and ran three lab replicates for each isolate in each of its test environments, with the exception of *Trichoderma koningii* for which we used three independent biological isolates per treatment for an initial proof-of-concept study. Each Petri dish was inoculated with one plug of an isolate’s stock culture placed on top of the litter at the center of each plate (Fig. S3a). Petri dishes were incubated for seven weeks at 25°C in darkness (Fig. S3b). At harvest, all litter was removed from each Petri dish by hand using forceps, weighed, and subsequently oven dried at 60°C for 48 hr to determine moisture content. The ability of each isolate to decay litter in each environment was calculated as percent litter mass loss (dry wt basis).

### Statistical analyses

Two-way analysis of variance (ANOVA) was used to test whether longterm exposure to N enrichment significantly affected the growth rate or litter mass loss associated with each species, with isolate origin and laboratory environment as independent variables (R version 3.0.1, R Core Team, 2013). We compared the responses of N50 or N150 isolates with control isolates of the same species. When interactions between isolate origin and laboratory environment were significant, we analyzed the data of each environment separately to test if isolates of different origin behaved differently in a common environment. When necessary, variables were transformed to ensure a normal distribution and homogeneity of variance. Using the same approach, we also tested for a general fungal response, by pooling the data across all species.

## RESULTS AND DISCUSSION

Global change experiments offer a unique opportunity to investigate and record evolutionary responses to rapid environmental change in the field, but their potential remains largely unexploited (Weese et al. 2015; Bataillon et al. 2016). Microbial evolution is likely to impact critical ecosystem processes, including organic matter decomposition. We tested whether chronic soil N enrichment has affected the ability of diverse fungi to decompose plant material. Using a common garden approach, we grew control and N-exposed fungi isolated from a 28 year simulated N deposition experiment in identical laboratory environments to test whether fungi exposed to chronic N enrichment consistently grow differently, or decompose litter differently, compared to control fungi. Experiments tested whether environment is the main driver of behaviors by comparing the same individuals across different laboratory environments, while simultaneously testing whether the identity of the individuals themselves is instead the main driver of behaviors by comparing different individuals isolated from different treatment plots (from control plots or N exposed plots) in the same laboratory environments (the same “gardens”). To our knowledge, this study is the first to test if and how fungal behaviors relevant to a critical ecosystem process evolve in response to long-term environmental change.

### Growth rates

All species grew with a single growth front and in a consistent linear fashion along race tubes, except *Cladosporium* whose growth was patchily distributed throughout the race tubes. This growth pattern made it impossible to calculate a growth rate and so this species was excluded from subsequent growth rate analyses. The remaining species grew at very different rates; the rapidly growing *Trichoderma* raced at ~22 mm dy^−1^ while growth rates of the other species ranged from 1.0 to 10 mm dy^−1^.

While biological replicates of a species from a specific treatment plot behaved similarly to each other, different species showed idiosyncratic growth responses, with one species *(Paraconiothyrium)* exhibiting faster growth of N isolates compared to control isolates and two species *(Cylindrium, Trichoderma)* showing slower growth (Fig. 1). Two species *(Alternaria, Discosia)* showed no response of long-term N enrichment on growth rate, while four species *(Epicoccum* I and II, *Pestalotiopsis, Phacidium)* showed mixed results. Thus the majority of ascomycete species proved sensitive to chronic N additions, with isolates from N amended plots growing differently (faster or slower) from control isolates, even when N isolates were grown in the away (control, low N) environment. However, the direction of change was not predictable, with the average response being no effect. The basidiomycete *Irpex* showed significantly higher growth rates for the N isolates (9.8–10 mm dy^−1^) in both their home (N150) and away (N0, control) environments compared to the control isolates (7.6–7.8 mm dy^−1^; origin: *P* < 0.0001, environment: *P* = 0.1976; Fig. 2A).

**FIG. 1.**
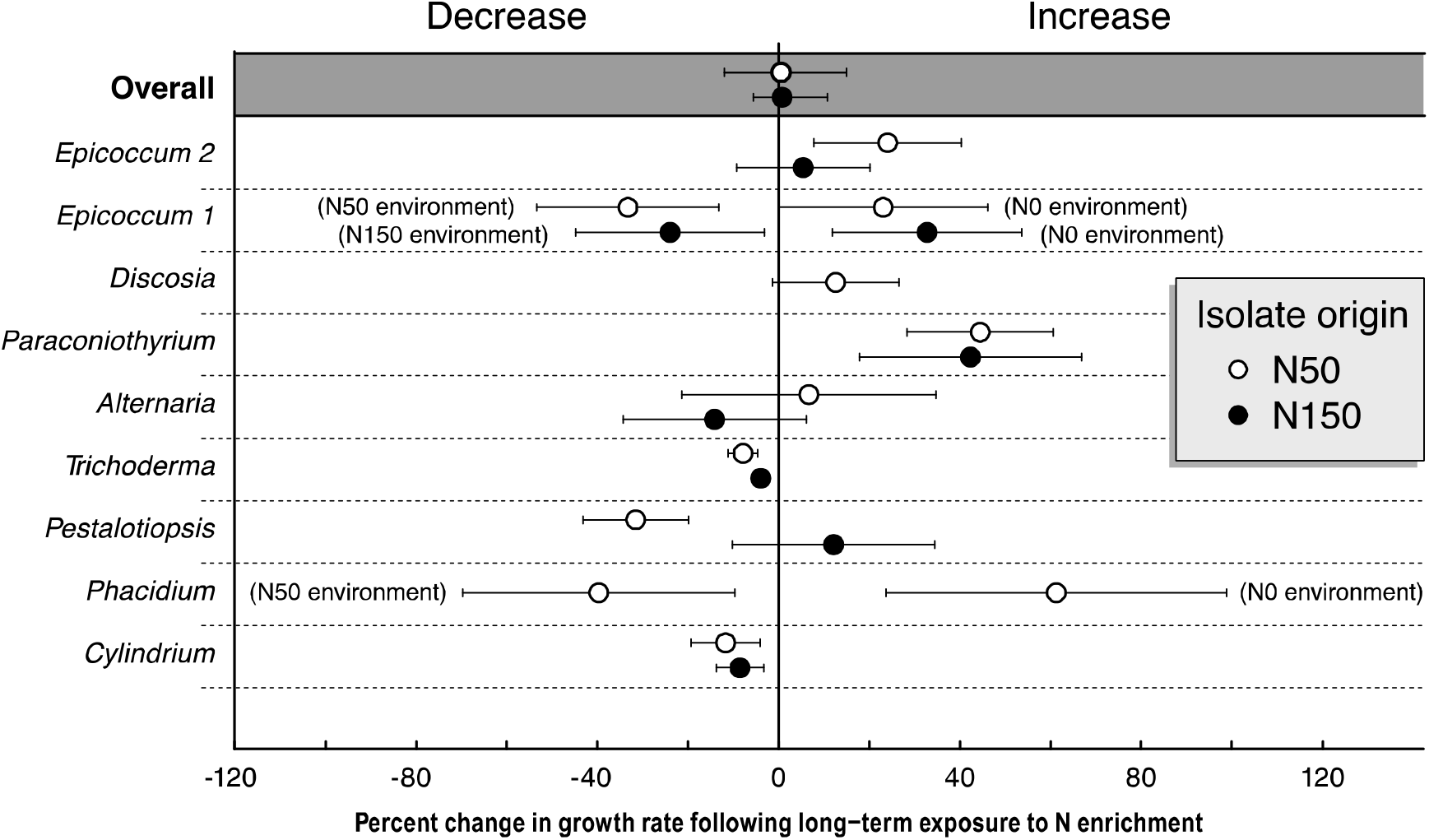
Percent change in growth rates of cellulolytic ascomycetes following long-term exposure to soil N enrichment. Open and filled circles represent N50 or N150 isolates, respectively, compared to control (N0) isolates. Note *Discosia* and *Phacidium* were only isolated from control plots and one of the two N treatment plots. Error bars are 95% confidence intervals. When error bars do not overlap with zero, the change in growth rate is significant at α < 0.05. Because there was a significant statistical interaction between isolate origin and laboratory (common garden) environment for *Epicoccum* 1 and *Phacidium*, the growth rates of these species in each of the lab environments are shown separately.

**FIG. 2.**
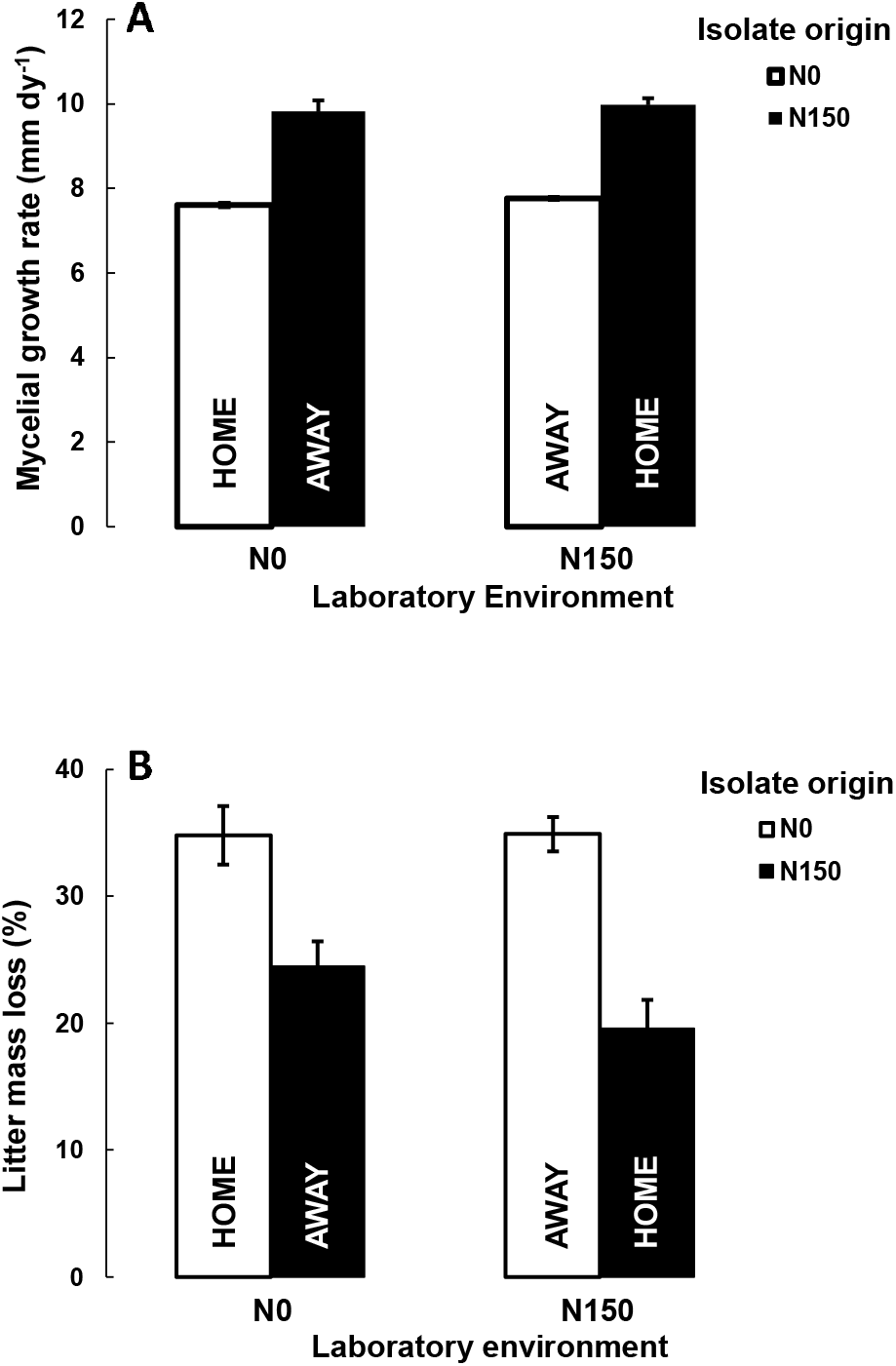
Mycelial growth rate (A) and litter mass loss (B) (mean and SE) for the white rot basidiomycete *Irpex* grown in home and away environments. Isolates of *Irpex* were not cultured from the N50 treatment.

### Litter decay

While growth responses were inconsistent, the capacity to decay plant litter, potentially a more relevant measure of ecosystem function, was significantly and consistently less for fungi isolated from N enriched plots compared to the same species isolated from the control treatment. Mass loss of litter averaged 19.6–34.9% for the white-rot fungus *Irpex*, compared to 7.0–12.6% for the ascomycete species (Figs. 2B and 3), which is not surprising since white-rot fungi can typically fully decompose lignocellulose, while most ascomycetes specialize in cellulose breakdown and only partially decompose lignin. Mass loss for litter decomposed by N isolates of *Irpex* was significantly lower compared to control isolates and this result was independent of the laboratory environment in which the isolates were grown (origin: *P* < 0.0001; environment: *P* = 0.3803; Fig. 2B). Mass loss for litter decomposed by ascomycete species was on average 17.9 and 26.2% lower for N50 and N150 isolates, respectively, compared to control isolates (Fig. 3). While isolate origin was significant (P < 0.01), there was no significant origin × environment interaction, meaning that N50 and N150 isolates decomposed litter more slowly than control (N0) isolates, irrespective of the laboratory environment in which they were grown (Fig. 3 inset). In terms of species-level responses, *Cylindrium* proved to be an exception and the only species able to decay litter more effectively after exposure to chronic N enrichment and this was only for isolates collected from the highest N addition (N150) treatment. There was no relationship between litter mass loss and growth rates (r^2^ = 0.04; *P* = 0.1063; Fig. S4), suggesting that growth rate, as measured here, is not a useful predictor of decay dynamics.

**FIG. 3.**
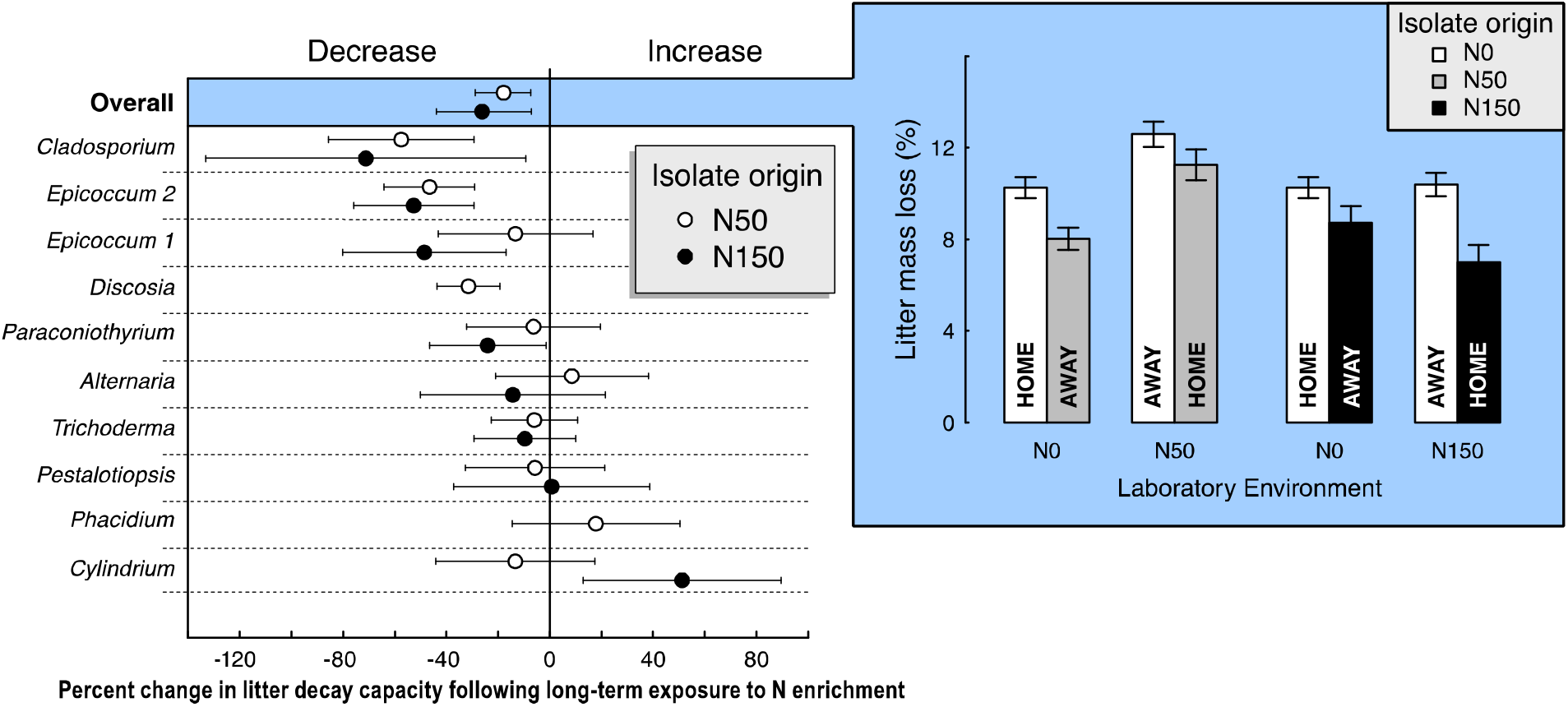
Percent change in litter decay capacities of cellulolytic ascomycetes following long-term exposure to soil N enrichment. Open and filled circles represent N50 or N150 isolates, respectively, compared to control (N0) isolates. Note *Discosia* and *Phacidium* were only isolated from control plots and one of the two N treatment plots. Error bars are 95% confidence intervals. When error bars do not overlap with zero, the change in litter mass loss is significant at a < 0.05. There were no significant interactions between isolate origin and laboratory environment. Inset graph on right shows litter mass loss averaged across all ascomycete species grown in their home versus away environment. Error bars indicate one standard error of the mean.

### Synthesis

In the aggregate, our data suggest that fungi growing in N enriched plots have evolved and are less able to decompose plant litter than the same species growing in control plots. Individual fungal isolates behaved consistently across home and away environments; an isolate less able to decompose in one environment was less able to decompose in all environments. Isolates from N treatment plots were generally less able to decay litter than isolates of the same species from control plots, and the behaviors of these N isolates did not revert or resemble control isolate behaviors even when the N isolates were grown in control environments. An especially striking difference was recorded among isolates of the basidiomycete, lignin-degrading species *Irpex;* litter mass loss was 30–44% lower for the N isolates compared to the control isolates in both control and N enriched laboratory environments.

Whether or how these altered behaviors will benefit a fungus remains untested and would require the tracking of entire life cycles in nature. The mechanism causing behavioral changes also remains unknown. Changes may be mediated by altered patterns of gene expression, including, for example, gene expression alterations caused by novel patterns of DNA methylation; changes in allele frequencies between the populations of control and N treatment plots so that rare alleles become dominant in treatment plots; novel DNA sequence evolution with downstream effects on enzyme functions; or some combination of these or additional forces. Regardless of mechanism, it appears that long-term exposure to chronic N additions may have acted as a novel selective pressure on the behaviors of these saprotrophic fungi. And while there may be gene flow across field treatment plots, the fact that we found consistent changes in species behaviors suggests that whatever dispersal there is among plots is not enough to counter natural selection. Our data connect to a classic literature on plant adaptation, including experiments documenting rapid adaptation to pollution and across sharp clines or adjacent treatment plots (e.g., Snaydon and Davies 1982). Early research on the heavy metal tolerance of plants growing on toxic mine spoils (Antonovics et al. 1971) served to both document plant adaptation to environmental change and provide evidence for genetic differences among closely spaced populations. The fungal behaviors described here are clearly an analogous result.

Previous research at the CNAS has demonstrated that N enrichment increases the species richness and diversity of ascomycetes generally (cellulose degraders), with an especially pronounced effect on the relative abundance of several ascomycete genera *(Trichophyton, Phialophora, Hypocrea/Trichoderma*, and *Rhizoscyphus* spp.) (Morrison et al. 2016). Others have also observed increased ascomycete richness and abundance with soil N enrichment (Allison et al. 2007; Weber et al. 2013). A shift in the fungal community toward taxa whose primary niche appears to be cellulose (rather than lignin) decay coupled with the fundamental changes in fungal behaviors we observed here (among ascomycetes and a basidiomycete) may accompany or perhaps even drive previously observed N-induced declines in fungal biomass, ligninolytic enzyme activity, and plant litter decay (Magill and Aber, 1998; Frey et al. 2004, 2014). Because observed changes in decay abilities were not readily reversed when N isolates were grown in control environments, whether or how quickly a fungal community will recover following the cessation of N enrichment emerges as another critical, unanswered question. Meanwhile, and for as long as N pollution continues to be a feature of our rapidly changing world, the evolution of fungal behaviors in these novel selective contexts will likely shape ecosystem function.

## ACKNOWLEDGMENTS

This work was supported by a National Science Foundation grant (DEB-1021063) to SDF and AP, a NSF Graduate Research Fellowship grant (DGE-1325256) to EAL, and by the NSF Long-Term Ecological Research (LTER) Program at Harvard Forest. Special thanks to Lisa Graichen, Mel Knorr, and Meghan Thornton for lab assistance.

## SUPPORTING INFORMATION

Additional supporting information may be found in the online version of this article at <u>http://onlinelibrary.wiley.com/</u>.

